# Histone H1 Variants Regulate Neurodevelopmental Transcriptional Programs in Autism with 16p11.2 deletion

**DOI:** 10.64898/2026.04.15.718677

**Authors:** Roni Brudno, Dan Askayo, Doha Khair, Ronna Shayevitch, Ifat Keydar, Michal Zmudjak-Olevson, Galit Lev-Maor, Mihaela Zavolan, Ran Elkon, Gil Ast

## Abstract

**Background:** Neurodevelopmental disorders, including autism spectrum disorder, involve widespread transcriptional dysregulation. Copy number variations at 16p11.2 are among the strongest genetic risk factors for autism spectrum disorder, yet the molecular mechanisms by which these copy number variations contribute to neurodevelopmental pathology remain unclear.

**Results:** We identify significant genetic associations between autism spectrum disorder susceptibility and the HIST1 histone gene cluster through genome-wide analysis. Transcriptomic profiling across post-mortem brain tissue, patient-derived neural progenitor cells, neurons, and cerebral organoids reveals consistent upregulation of linker histone variants H1.2 and H1.5 in idiopathic autism spectrum disorder and 16p11.2 hemi-deletion carriers, but not in schizophrenia or bipolar disorder. Functional assays demonstrate that dysregulated H1 expression disrupts gene networks involved in synaptic signaling, chromatin remodeling, and neural differentiation. Mechanistically, we link H1 upregulation to MAZ, a transcription factor encoded within the 16p11.2 locus. MAZ binds the promoter regions of H1 genes and represses their transcription. Knockdown of MAZ leads to H1 overexpression. H1 upregulation alone is sufficient to alter the expression of autism spectrum disorder-associated genes.

**Conclusions:** Our findings define a MAZ-dependent regulation of H1 dosage as a critical chromatin-mediated mechanism contributing to transcriptional pathology in 16p11.2-associated autism spectrum disorder.

## Background

Neurodevelopmental disorders such as autism spectrum disorder (ASD) have complex genetic architectures, with hundreds of risk variants identified across the genome [1]. The majority of these associated variants reside in noncoding or regulatory regions, implicating disruption of gene expression regulation as a primary mechanism in disease pathogenesis [2]. Consistently, large-scale transcriptome studies of post-mortem brains have revealed widespread dysregulation of gene expression and mRNA splicing in ASD, affecting a substantial fraction of the transcriptome and converging on neuronal and immune pathways [3–5]. These observations point to transcriptional regulatory mechanisms as key mediators of ASD risk.

ASD affects approximately 1% of the population and exhibits strong heritability, with a 10-fold increased risk in first-degree relatives [6]. Yet genetic diagnosis remains limited due to incomplete penetrance, as ASD-associated variants are frequently carried by unaffected individuals [7]. This “missing heritability” suggests that additional factors beyond inherited variants may influence disease manifestation, particularly through altered gene expression patterns and chromatin-mediated regulation [8]. A key example is the deletion of a segment on one of the two copies of chromosome 16. This is known as a 16p11.2 hemi-deletion, and it affects a region containing 29 genes, of which 27 are protein-coding genes involved in synaptic function, transcriptional regulation, and chromatin remodeling—core processes for brain development [9, 10]. The 16p11.2 hemi-deletion represents one of the strongest genetic risk factors for ASD, yet exhibits only 20–30% penetrance in carriers [11, 12]. These observations highlight the need to understand how gene-regulatory networks and chromatin context modulate risk conferred by high-impact variants such as 16p11.2. Central to this question are the regulatory drivers located within the deletion itself; among the 27 protein-coding genes in the region, there are two sequence-specific transcription factors: T-box transcription factor 6 (TBX6) and MYC-associated zinc finger protein (MAZ) (Fig. 1A).

**Figure 1.**
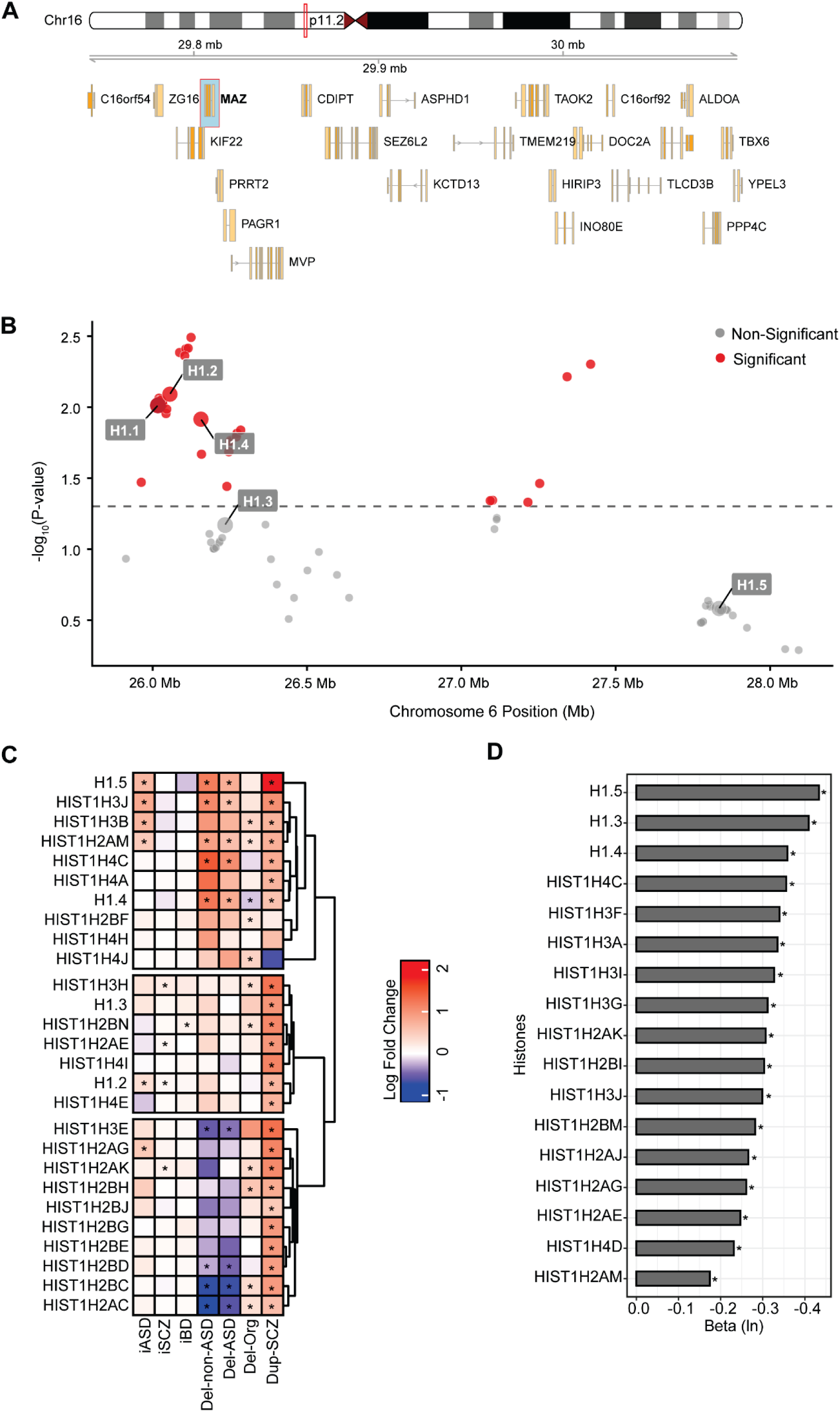
Gene-scoring and Expression patterns of the HIST1 gene cluster in idiopathic and 16p11.2 CNV-associated ASD. (**A**) Genomic map of the MAZ-containing sub-interval within the 16p11.2 copy number variant (chr16: 29,700,000–30,097,000; GRCh38). The diagram displays the spatial organization of MAZ relative to its immediate flanking genes in the deletion region. (**B**) Manhattan plot showing gene-based associations of HIST1 cluster genes with ASD, computed using PascalX. Only genes within the HIST1 cluster on chromosome 6 are shown. Red markers indicate nominally significant associations (p < 0.05). (**C)** K-means clustering and heatmap of differentially expressed HIST1 genes (log₂ fold change; FDR-adjusted p < 0.05) across post-mortem brain tissue from individuals with idiopathic ASD (iASD), schizophrenia (iSCZ), and bipolar disorder (iBD), together with *in vitro* models including NPCs derived from individuals with 16p11.2 hemi-deletion (ASD [Del-ASD] and neurotypical [Del-non-ASD]), corresponding brain organoids derived from neurotypical NPCs (Del-Org), and NPCs from individuals with 16p11.2 duplication and SCZ (Dup-SCZ). (**D**) Correlation between HIST1 gene expression and 16p11.2 copy number variation in lymphoblastoid cell lines from ASD patients with 16p11.2 deletions or duplications. Data represents the natural logarithm of the beta coefficient derived from a linear regression modeling the effect of CNV dosage on gene expression (p < 0.05).

Histone proteins are central to chromatin organization and regulation of gene expression. In addition to the four core histones that form the nucleosome, the linker histone H1 binds at nucleosomal entry and exit sites to modulate higher-order chromatin structure and DNA accessibility [13]. The H1 family is the most divergent class of histones, comprising multiple variants in humans, including several somatic subtypes (H1.1–H1.5) with distinct temporal and tissue-specific expression patterns [14, 15]. They also contribute to transcriptional regulation, with H1.2 and H1.5 showing occupancy patterns that correlate with gene expression states [15, 16].

Links between H1 variants and human neurodevelopmental disease have only recently begun to emerge. A *de novo* frameshift mutation in *HIST1H1E* (encoding histone H1.4) was reported in a child with ASD and intellectual disability [17], and H1.2 (*HIST1H1C*) is significantly upregulated in post-mortem brain tissue from individuals with ASD [18]. H1.2 is the most abundant linker histone in human somatic cells and a key regulator of gene expression [15, 19]. Notably, the genes encoding H1.2 and H1.5 reside within the HIST1 histone supercluster in the major histocompatibility complex (MHC/HLA) region on chromosome 6, a genomic locus enriched for regulatory variation and expression quantitative trait loci [20–32]. Thus, both their biochemical functions and genomic context suggest that H1 variants in general, and H1.2 and H1.5 in particular, could play an important role in ASD-associated transcriptional dysregulation.

Here, we identify the potential role of histone H1s upregulation in the development of ASD. We first identified elevated expression of H1.2 and H1.5 in idiopathic ASD and in ASD cases carrying the 16p11.2 hemi-deletion. We demonstrate that overexpression of these H1 variants, without the 16p11.2 copy number variation (CNV), perturbs gene expression networks involved in synaptic signaling, chromatin remodeling, and neural differentiation. Importantly, we uncover a mechanistic link between 16p11.2 and the HIST1 cluster through the transcription factor MAZ, encoded within the 16p11.2 region. We show that MAZ acts as a repressor at H1 gene promoters, and that knockdown of MAZ expression leads to H1 overexpression. Together, these findings implicate MAZ-dependent control of H1.2 and H1.5 as a plausible chromatin-mediated mechanism linking 16p11.2 dosage to widespread transcriptional dysregulation in ASD.

## RESULTS

### Histone H1s Are Associated with ASD and Upregulated in Idiopathic ASD and 16p11.2 CNV Conditions

Given the growing evidence implicating chromatin regulation and epigenetic modifications in developmental disorders, we first asked whether histone genes within the HIST1 locus are associated with ASD. Using publicly available ASD genome-wide association studies (GWAS), including a meta-analysis of European ancestry samples, we performed a gene-based association test on 1050 genes from chromosome 6, focusing on the HIST1 cluster. Within this region, H1.2 (*HIST1H1C*) showed the strongest gene-based association signal, achieving nominal significance (p < 0.05). Similarly, other linker histones, including H1.1, H1.3, and H1.4, also displayed nominally significant associations (Fig. 1B). These results indicate that common variation in specific HIST1 genes, specifically H1.2, the most highly expressed somatic histone, may contribute to ASD susceptibility.

To further examine the roles of HIST1 genes in the etiology of neurodevelopmental disorders, primarily those associated with 16p11.2 copy number variations (CNVs), we utilized a comprehensive set of RNA-seq datasets. These included brain samples from individuals diagnosed with idiopathic ASD (iASD), SCZ (iSCZ), and BD (iBD). Additionally, we incorporated gene expression data from lymphoblastoid cell lines (LCLs) derived from individuals from ASD-affected families with the 16p11.2 CNVs, as well as RNA-seq datasets from iPSC-derived neural progenitor cells (NPCs) from individuals with 16p11.2 hemi-deletion with and without ASD (Del-ASD; Del-non-ASD, respectively), cerebral organoids generated from 16p11.2 hemi-deletion iPSC lines (Del-Org), and NPCs from individuals with 16p11.2 heterozygous duplication and SCZ (Dup-SCZ) (Fig. 1C). LCLs were used as a complementary peripheral model of 16p11.2 dosage and interpreted in parallel but separately from neural samples.

By focusing on the HIST1 gene cluster, we analyzed overlapping genes across multiple RNA-seq datasets and identified distinct expression patterns under specific conditions. In idiopathic ASD (iASD), several HIST1 genes—including H1.5 (*HIST1H1B*), H3C12 (*HIST1H3J*), H3C2 (*HIST1H3B*), H2AC17 (*HIST1H2AM*), H2AC11 (*HIST1H2AG*), and H1.2 (*HIST1H1C*)—showed a statistically significant increase in expression compared to idiopathic SCZ (iSCZ) and idiopathic BD (iBD) (Fig. 1C). Notably, H1.5 was consistently upregulated across iASD and multiple 16p11.2 CNV-associated conditions, suggesting a prominent role for this variant as a driver of transcriptional changes in these disorders.

We further observed markedly enhanced expression of H1.5, HIST1H3J, HIST1H2AM, and H1.2 in NPCs derived from individuals with 16p11.2 hemi-deletions and duplications. In contrast, these same HIST1 variants showed reduced or unchanged expression levels in iSCZ, iBD, and 16p11.2 hemi-deletion organoids (Del-Org) (Fig. 1C). This differential pattern indicates a strong context dependence of HIST1 dysregulation, with the most pronounced changes occurring in early neural progenitors and in the presence of 16p11.2 CNVs. These findings are in line with previous observations that both genetic background and developmental stage can profoundly influence histone gene expression dynamics [16, 25, 33].

Among the HIST1 genes, H1.5 exhibited the most significant and consistent elevation across iASD and 16p11.2 CNV-associated conditions. When we examined the effect of 16p11.2 dosage in LCLs, H1.5 displayed a significant negative beta coefficient (Fig. 1D). This indicates a negative correlation between copy number and expression: deletion carriers exhibit upregulation, while duplication carriers show reduced expression. This negative dosage effect suggests that the 16p11.2 locus harbors a negative regulator of H1.5, where the loss of this repressor in the deletion context leads to the observed upregulation. H1.2 also displayed significant, albeit more moderate, elevation across iASD, iSCZ, and Dup-SCZ. Importantly, H1.2 is the most abundant linker histone in human somatic cells and a key regulator of transcription, and both H1.2 and H1.5 show occupancy patterns that decrease from poorly to highly expressed genes [14, 34]. Upregulation of H1.2 in ASD brain tissue has been reported previously [17], and is recapitulated in our analysis.

Our findings suggest the involvement of H1.2 and H1.5 in the molecular pathophysiology of ASD. The coordinated elevation of H1.2 and H1.5 across independent datasets suggests a shared regulatory control within the HIST1 cluster, potentially involving cross-regulatory mechanisms between H1 variants as demonstrated by Pascal et al. [15]. These results point to a mechanistic framework in which dysregulation of H1.2 and H1.5 contributes to genome-wide transcriptional changes characteristic of ASD in general and 16p11.2 hemi-deletion more specifically.

### Transcriptional Landscapes Across Neurodevelopmental Stages

To comprehensively investigate the downstream effects of histone H1 variant dysregulation, we overexpressed (OE) and knocked down (KD) the H1.2 and H1.5 variants, either individually or in combination, in neuroblastoma (SH-SY5Y) cell lines and performed RNA-seq for each condition. We confirmed the efficiency of the overexpression and siRNA-mediated knockdown using qPCR and Western blot (Additional file 1: Fig. S1–S3). We further analyzed RNA-seq data previously generated by us from embryonic kidney (HEK293) cell lines in which somatic histone H1 variants were knocked out (KO) [15]. These cell lines include: 2KO, a double KO lacking H1.2 and H1.3; 4KO, a quadruple KO lacking H1.0, H1.2, H1.3 and H1.5; and 5KO, a quintuple KO lacking H1.0, H1.2, H1.3, H1.4 and H1.5.

These datasets were integrated with RNA-seq from iPSC-derived NPCs from individuals with 16p11.2 hemi-deletions, both with and without ASD (ASD-NPC and Non-ASD-NPC, respectively), cerebral organoids generated from 16p11.2 hemi-deletion iPSC lines without ASD (Organoids), NPCs from individuals with 16p11.2 duplication and SCZ (SCZ-NPC), and neurons from individuals with 16p11.2 duplication and SCZ (SCZ-neurons; Parnell et al., 2023) [35].

Using principal component analysis (PCA) and correlation approaches, we examined the transcriptional profiles of the conditions above. We observed a neurodevelopmental pattern, spanning NPCs to differentiated neurons and cerebral organoids, as indicated by a trend line (Fig. 2A).

**Figure 2.**
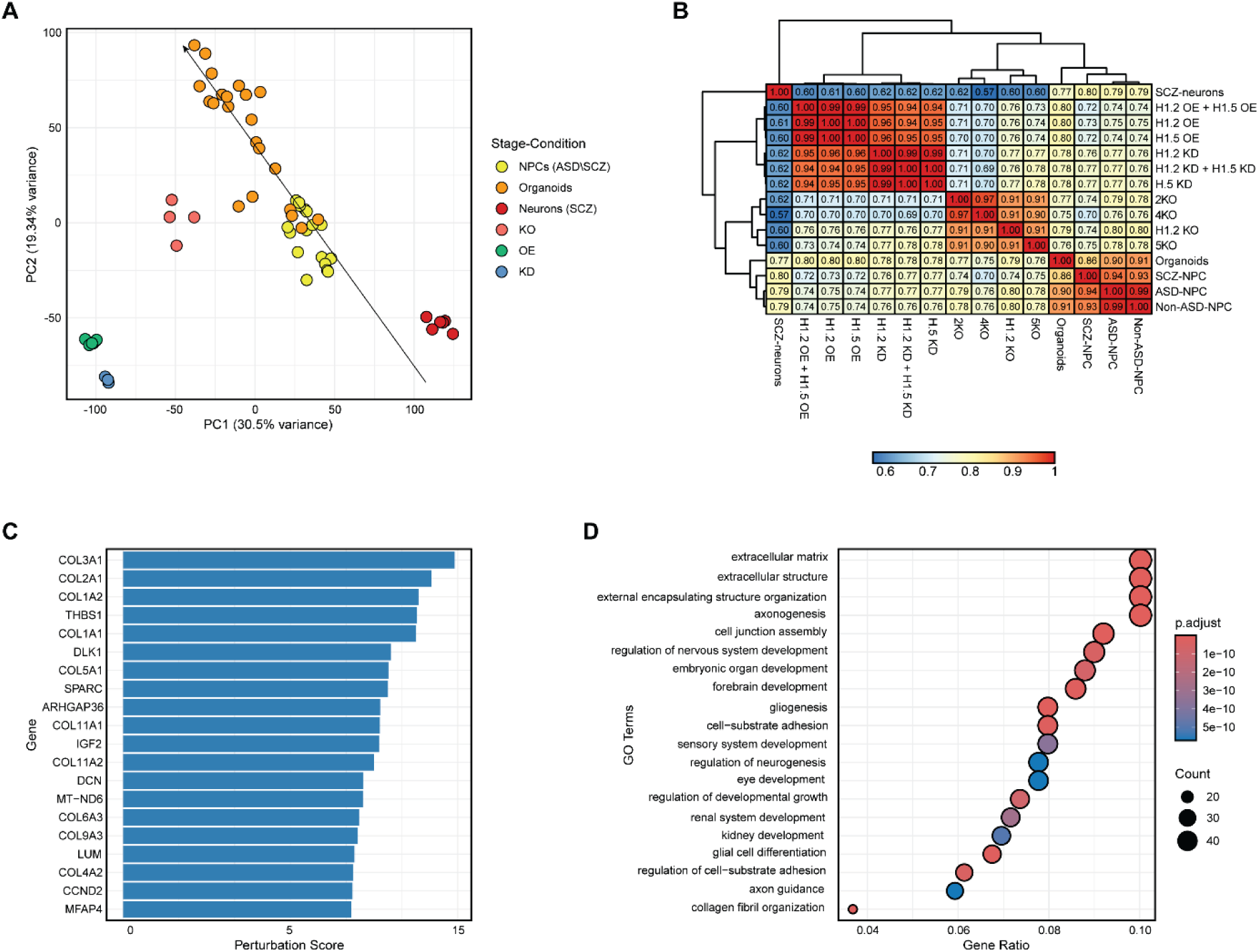
Transcriptional Landscapes of 16p11.2 Hemi-Deletion, Duplication, and Linker Histone Perturbation Models of Neurodevelopment. (**A**) PCA plot illustrating global transcriptional patterns across multiple experimental paradigms: NPCs harboring 16p11.2 hemi-deletion from both ASD (ASD-NPC) and neurotypical individuals (Non-ASD-NPC); cerebral organoids differentiated from neurotypical NPCs (Organoids); NPCs and mature neurons derived from individuals with 16p11.2 duplication and schizophrenia (SCZ-NPC, SCZ-Neurons); and SH-SY5Y neuroblastoma cells with genetic manipulations of linker histones H1.2 and H1.5 (overexpression (H1.2 OE, H1.5 OE, H1.2 OE & H1.5 OE), knockdown (H1.2 KD, H1.5 KD, H1.2 KD & H1.5 KD), and knockout (2KO, 4KO, 5KO, H1.2 KO)). PC1 and PC2 axes show the percentage of variance explained by each component. (**B**) Pearson correlation matrix displaying the pairwise relationships between gene expression profiles across all experimental conditions. Color intensity represents correlation strength. Hierarchical clustering was applied to groups with similar expression patterns. (**C**) Ranked bar plot highlighting the top 20 genes showing the largest transcriptional variability across conditions. Perturbation scores were calculated as the standard deviation of normalized gene expression levels across all models, reflecting the magnitude of expression change relative to matched control conditions. (**D**) Gene Ontology enrichment analysis of the 500 most perturbed genes. Dot size represents the number of genes in each GO term category, while color intensity indicates statistical significance (-log_10_ adjusted p-value). Only biological processes with FDR < 0.05 are shown.

Focusing on NPCs, we observed similar transcriptional profiles among individuals with 16p11.2 hemi-deletions, regardless of ASD diagnosis, and those with 16p11.2 duplications associated with SCZ. This unexpected similarity suggests that early neurodevelopmental transcriptional mechanisms remain remarkably consistent, despite underlying genetic differences (Fig. 2A, see Discussion). At later differentiation stages, 16p11.2 hemi-deletion brain organoids and SCZ-associated neurons with the 16p11.2 duplication diverged, revealing distinct neurodevelopmental trajectories (Fig. 2A-B). This shift underscores the dynamic nature of gene expression during neural development. Interestingly, histone variants H1.2 and H1.5 displayed a distinct pattern: OE and KD clustered together, separate from all other samples, whereas KO clustered closer to brain organoids (Fig. 2A). This suggests that altering H1.2 and H1.5 levels through OE or KD results in one transcriptional state, whereas complete KO leads to a fundamentally different profile.

To determine which experimental perturbation best recapitulates the 16p11.2 transcriptional state, we performed overlap analysis and PCA clustering of differentially expressed genes (Additional file 1: Fig. S4). We found that histone OE and KD conditions cluster closely with 16p11.2 hemi-deletion NPCs and organoids, sharing a significant intersection of regulated genes. This contrasts with complete knockout (KO), which induces a distinct profile, demonstrating that the partial H1 dosage imbalance modeled by OE/KD recapitulates key transcriptional features of the 16p11.2 hemi-deletion (both ASD and non-ASD lines).

To identify the specific genes that are most susceptible to histone H1 dosage imbalance, irrespective of the specific perturbation direction, we ranked candidates by their cross-condition variability (Fig. 2C). Top candidates included COL1A1, SPARC, and DLK1. To determine the biological impact of these dosage-sensitive changes, we performed enrichment analysis, which revealed that these dysregulated genes are heavily enriched in processes controlling brain development. The most affected pathways included the organization of the extracellular matrix, axonogenesis, cell junction assembly, and forebrain development (Fig. 2D). These results highlight the significant impact of histone variant dysregulation on neural differentiation programs, supporting previous observations [36, 37].

The similar gene patterns we observed in NPCs, which diverge as the cells mature into neurons, suggest that H1.2 and H1.5 are key regulators of early brain development. Our findings show that increases or decreases in H1 levels and not complete loss can shift gene expression programs in a way that resembles the 16p11.2 hemi-deletion state. These early changes may initiate a common molecular trajectory in ASD and SCZ, with disease-specific features emerging as cells progress through neuronal differentiation.

### Dysregulated Gene Expression Within the 16p11.2 Chromosomal Region and ASD-Associated Genes

To deepen our understanding of histone dysregulation impact on neurodevelopmental disorders, we conducted a focused analysis of gene expression within the 16p11.2 chromosomal region. We evaluated the specific contribution of individual histone variants by examining transcriptional profiles of H1.2 and H1.5 perturbations separately. The conditions included H1.2 OE, H1.2 KD, H1.5 OE, H1.5 KD, and combined perturbations (Both OE, Both KD).

Expression patterns of 16p11.2 genes revealed a clear distinction based on dosage. Combined overexpression of H1.2 and H1.5 most closely matched the downregulation profile seen in 16p11.2 hemi-deletion models (ASD, Non-ASD, and Organoids). In contrast, single overexpression of either H1.2 or H1.5 more closely resembled the upregulation observed in duplication-associated SCZ models (Fig. 3A). This bidirectional behavior suggests that H1.2 and H1.5 can act together to shift the transcriptional state toward a “deletion-like” profile, while individual overexpression tends to promote a “duplication-like” expression pattern.

**Figure 3.**
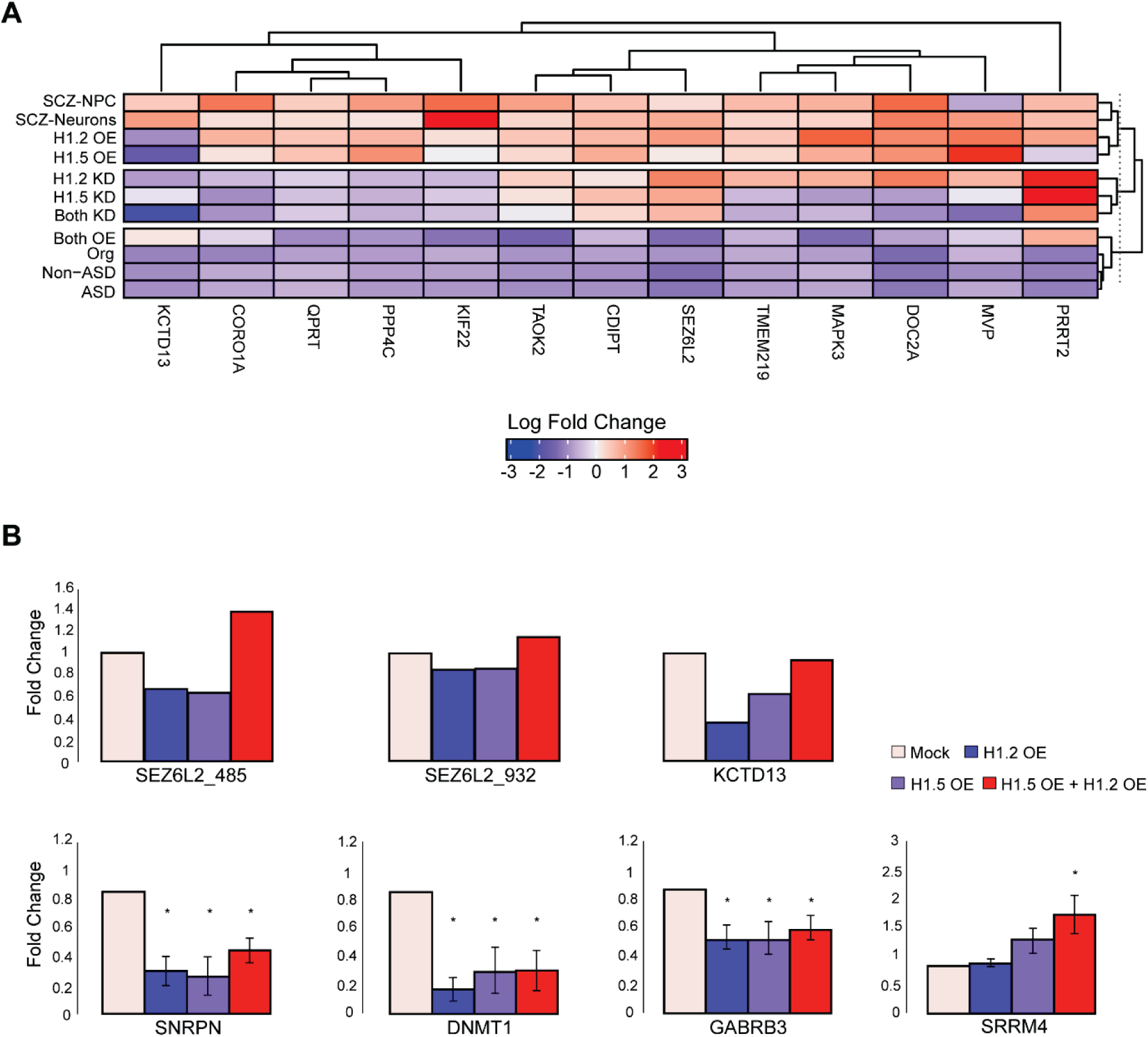
Expression Profiles of 16p11.2 Region Genes and ASD-associated genes. (**A**) Heatmap showing fold-change values for significantly differentially expressed genes within the 16p11.2 region across histone H1 perturbation conditions, relative to matched control samples. Histone perturbations are shown separately for H1.2 overexpression (H1.2 OE), H1.2 knockdown (H1.2 KD), H1.5 overexpression (H1.5 OE), H1.5 knockdown (H1.5 KD), and combined manipulation of both histones (Both OE, Both KD). (**B**) RT-qPCR analysis of mRNA levels. Expression of 16p11.2 loci gene isoforms (SEZ6L2_485, SEZ6L2_932, *KCTD13*, n = 1) and ASD-associated genes (*SNRPN*, *DNMT1*, *GABRB3*, *SRRM4*, n = 3) in cells overexpressing H1.2 (H1.2 OE), H1.5 (H1.5 OE), or both histones (H1.2 OE & H1.5 OE). Mock (empty vector) was used as the control. Data shown as mean ± SEM. *p < 0.05.

To further test this dynamic effect, we performed RT-qPCR analysis on selected 16p11.2 and ASD-associated genes (Fig. 3B). Consistent with the global profiling, the combined overexpression of H1.2 and H1.5 resulted in a distinct expression profile compared to the single overexpression conditions, confirming that the simultaneous elevation of both variants drives a qualitative shift in gene regulation.

Our findings suggest that these histone variants function as critical regulatory elements capable of modulating gene expression in different genome parts. Importantly, these histones play a significant role in controlling gene activity in ASD, particularly in the 16p11.2 region associated with these neurodevelopmental disorders.

### Functional Network Analysis of Dysregulated Pathways

To identify molecular signatures shared across distinct genetic contexts of neurodevelopmental disorders, we performed a protein-protein interaction (PPI) network analysis based on intersecting differentially expressed genes (DEGs). Genes were included only if they were significantly differentially expressed (FDR < 0.05) relative to their matched controls in all of the following conditions: brain organoids derived from 16p11.2 hemi-deletion models (Org), neurons from individuals with 16p11.2 duplication and schizophrenia (SCZ-Neurons), H1.2 overexpression (H1.2 OE), H1.5 overexpression (H1.5 OE), and combined H1.2/H1.5 overexpression (Both OE). This stringent intersection defines a core set of genes that are consistently dysregulated across both histone H1 dosage perturbations and 16p11.2 CNV-associated neural models, including both upregulated and downregulated genes. Consistent with the presence of shared and context-specific transcriptional programs, an independent genome-wide analysis using a variational autoencoder identified histone-driven gene clusters. These clusters distinguish shared versus disorder-specific regulatory patterns across ASD, SCZ, 16p11.2 CNV, and H1 overexpression models (Additional file 1: Fig. S5).

The network revealed two primary modules: one associated with glutamatergic synapse function, and the other with the assembly of free 40S ribosomal subunits (Fig. 4A). The glutamatergic synapse cluster signifies the importance of synaptic signaling in ASD pathophysiology, as glutamatergic neurotransmission has been implicated as a key mechanism in this disorder [34, 38]. Genes within this cluster are directly linked to neuronal excitability and synaptic plasticity, processes critical for learning, memory, and cognitive function, which are often disrupted in neurodevelopmental disorders [39, 40]. The second cluster, focused on 40S ribosomal subunit assembly, highlights the role of translational control in neurodevelopmental disorders, an area of increasing interest for its contributions to synaptic and neuronal function [41, 42]. Dysregulation of ribosomal protein expression, as observed in this cluster, may reflect broader disruptions in protein synthesis pathways that disrupt neuronal development and connectivity.

**Figure 4.**
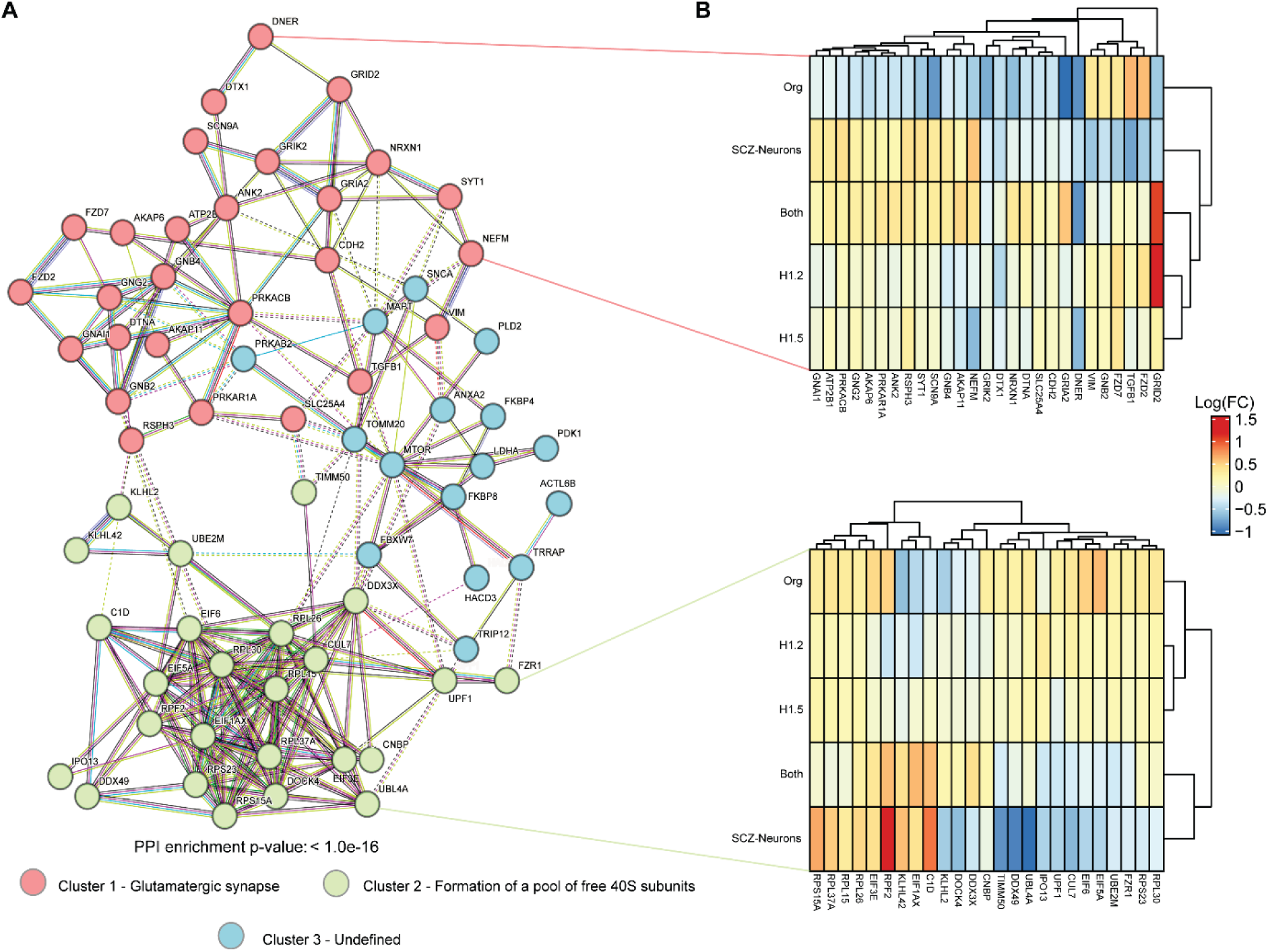
Protein–Protein Interaction Network Analysis of H1 Variant Overexpression and 16p11.2 Copy Number Variation Models. (**A**) PPI network of intersecting DEGs, defined as genes significantly differentially expressed (FDR < 0.05, relative to matched controls) in all of the following conditions: Org (brain organoids from 16p11.2 hemi-deletion models), SCZ-Neurons (neurons from individuals with 16p11.2 duplications and SCZ), H1.2 OE, H1.5 OE, and combined H1.2/H1.5 OE (Both OE). Nodes represent proteins and edges represent validated protein–protein interactions. The network separates into three modules: Cluster 1 enriched for glutamatergic synapse components, Cluster 2 enriched for 40S ribosomal subunit assembly, and a smaller Cluster 3 with heterogeneous or undefined functions. The network shows significant interaction enrichment (PPI enrichment p-value < 1.0 × 10⁻¹⁶). (**B**) Heatmap representation of expression changes (log_2_ fold change) for genes within the identified clusters across different experimental conditions. Dendrograms show clustering of genes and conditions based on expression similarity.

Gene expression analysis within these clusters demonstrated that the OE of both histones H1.2 and H1.5 had a pronounced effect, particularly within the glutamatergic synapse cluster. Transcriptional changes observed in this cluster mirrored those found in SCZ-associated neurons and 16p11.2 hemi-deletion-associated brain organoids (Fig. 4B). This alignment suggests that histone OE can recapitulate transcriptional disruptions across genetic contexts, further supporting its role as a driver of shared molecular mechanisms in ASD.

Together, these results highlight the potential of targeting histone-mediated pathways to address shared and distinct features of neurodevelopmental disorders.

### MAZ Links 16p11.2 Hemi-Deletion to Linker Histone H1 Dysregulation

To uncover the molecular mechanism connecting the 16p11.2 hemi-deletion to the elevated expression of H1 genes, we examined the functional characteristics of the 27 protein-coding genes within the region. We focused on the two sequence-specific transcription factors: TBX6 and MAZ. While TBX6 is restricted to paraxial mesoderm differentiation and linked to congenital scoliosis [43, 44], MAZ is ubiquitously expressed in the developing brain and functions as a dosage-sensitive regulator of GC-rich promoters [45, 46].

Motif analysis of the H1 cluster revealed that putative MAZ binding sites are significantly enriched near the transcription start sites (TSS; Fig. 5A). This architecture aligns with the intrinsic binding properties of MAZ, which globally favors GC-rich sequences (Additional file 1: Fig. S6A). As H1 promoters are similarly GC-rich, we prioritized MAZ as the key mediator connecting the 16p11.2 hemi-deletion to H1 transcriptional changes.

**Figure 5.**
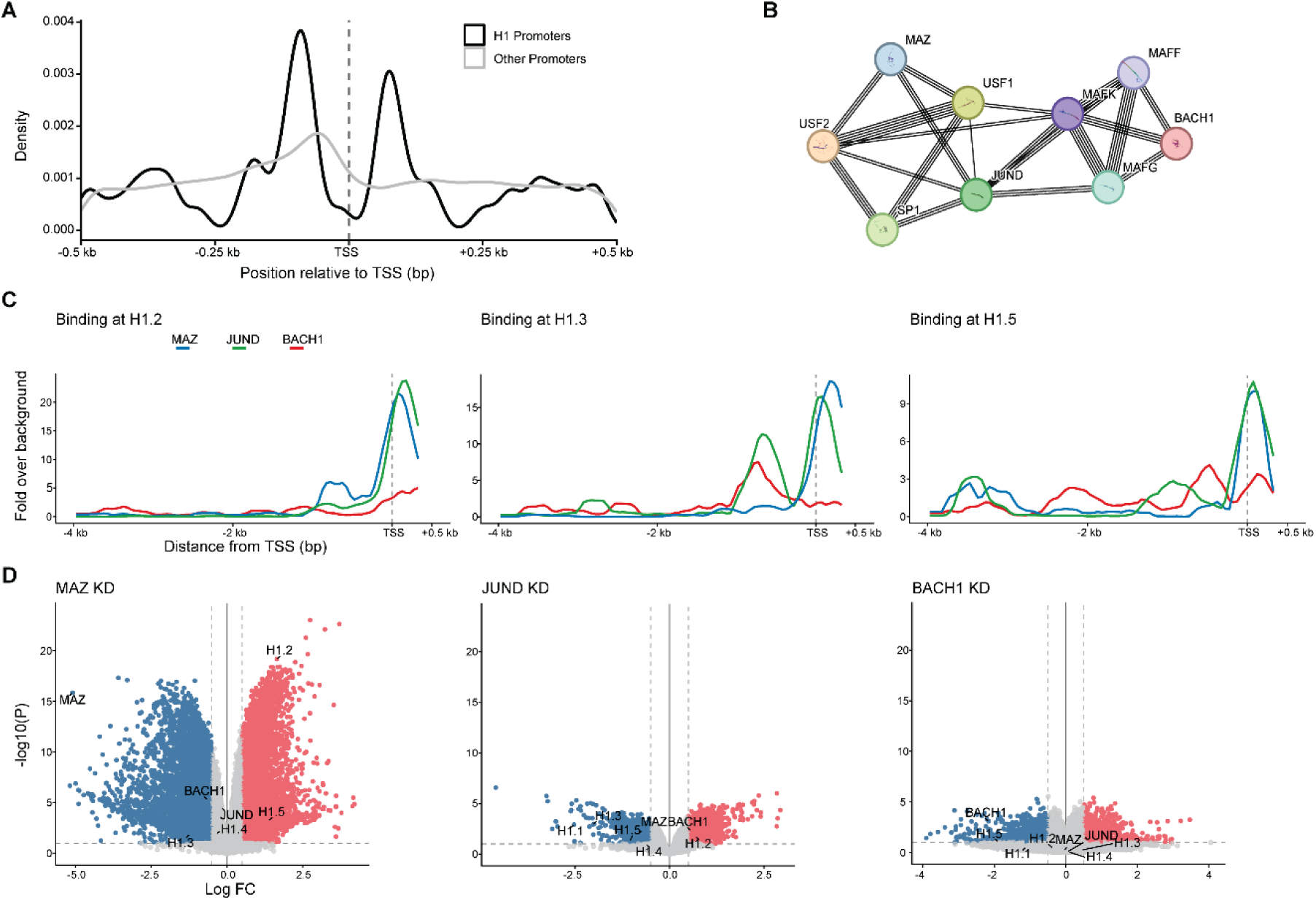
MAZ-centered regulatory network linking the 16p11.2 CNV to histone H1 promoter regulation. (**A**) Motif density profile showing the positioning of MAZ binding sites relative to the transcription start site (TSS). The distribution of MAZ motifs in H1 promoters (black line, n=61) is compared to a background of 5,000 random promoters (gray line), within a ±500 bp window relative to the TSS. (**B**) Protein–protein interaction network derived from STRING [47] analysis. The graph visualizes high-confidence physical and functional associations (combined score ≥ 0.4) between MAZ and other promoter-associated transcription factors, including JUND and BACH1. (**C**) Analysis of ChIP-seq data showing distinct binding profiles for MAZ, JUND, and BACH1 at the indicated genes. Curves show average signal expressed as fold over distal background, with position on the x-axis indicating distance from the TSS (vertical dashed line). (D) Volcano plots showing transcriptional changes following siRNA-mediated knockdown of MAZ, JUND, or BACH1 compared with nontargeting control siRNA. Each point represents a gene, with color indicating classification based on nominal p-value and effect size: upregulated (p < 0.1 and log₂ fold change ≥ 0.5), downregulated (p < 0.1 and log₂ fold change ≤ −0.5), or not significant. Horizontal dashed lines mark the p-value threshold (−log₁₀ p = 1), and vertical dashed lines indicate log₂ fold-change thresholds (±0.5). A common y-axis range is used across panels to facilitate comparison of effect sizes. Histone H1 genes and selected transcription factors from the MAZ network are labeled.

We first placed MAZ within a broader functional context, and we constructed a protein–protein interaction network using high-confidence data from STRING [47]. This analysis identified a tightly connected transcriptional module comprising MAZ and several key promoter-associated factors (Fig. 5B). We focused our attention on JUND and BACH1 due to the following biological reasons: JUND is established to cooperate with Sp1-like factors to drive transcription at GC-rich promoters [48], while BACH1 preferentially occupies promoters and enhancer-promoter hubs of highly active genes in embryonic stem cells, which are typically CpG-rich [49]. The robust connectivity of this module suggests that MAZ does not function alone but rather acts as a complex within a transcription factor hub that controls activity at GC-rich target sites.

We characterized the physical interactions between MAZ, JUND, and BACH1 across the TSS region of the different H1 genes expressed in somatic human cells (*HIST1H1A*–*HIST1H1E*; H1.1–H1.5). For that, we analyzed ChIP-seq data generated from K562 cells. Mapping binding enrichment across the TSS of H1 genes revealed distinct occupancy patterns. Both MAZ and JUND showed robust, broad enrichment across all the examined H1 genes, with peaks centered around the TSS (Fig. 5C and Additional file 1: Fig. S6B). H1.2 shows an additional minor binding site around 1 kb upstream from the TSS, distinguishing its upstream regulation from the other H1s. H1.3 also shows a unique binding profile with a binding of both JUND and BACH1 around 1 kb upstream of the TSS. Finally, H1.5 displayed several secondary binding sites upstream of the TSS of MAZ, BACH1, and JUND (one location of the secondary binding site of BACH1 is shared with H1.3). Extending this analysis to the full set of H1 variants showed that H1.4 exhibits a MAZ promoter peak similar to those shown in Fig. 5C, whereas H1.1 was the only H1 gene lacking a comparable MAZ enrichment at its promoter (Additional file 1: Fig. S6B). However, this is likely due to the very low transcription level of H1.1 in this cell line (data not shown). These results indicate that while MAZ localizes to the promoter regions of all examined H1 genes (except H1.1), its specific co-occupancy with JUND and BACH1 creates unique regulatory signatures for individual variants, supporting a model of precise combinatorial control (see Discussion).

To examine the functionality of these transcription factors on H1 transcription, we analyzed transcriptomic data from KD analysis of MAZ, JUND, and BACH1. Volcano plot analysis revealed that MAZ depletion induced a robust effect on the transcription levels of many genes, with a strong upregulation effect on multiple H1 genes, particularly H1.2 and H1.5 (Fig. 5D). This directional change indicates that MAZ functions primarily as a transcriptional repressor at these loci. In contrast, KD of its network partners produced distinct, gene-specific effects: JUND knockdown led to upregulation of H1.2 but downregulation of H1.5, while BACH1 knockdown led to the specific downregulation of H1.5 (Fig. 5D). Furthermore, silencing of SP1 (data not shown) resulted in downregulation restricted solely to H1.2.

To further assess the functional impact of these shifts on chromatin organization in SH-SY5Y cells, we integrated our transcriptomic data with ATAC-seq–derived maps of open chromatin. High-confidence open promoters were defined and evaluated using Gene Set Enrichment Analysis (GSEA) [50]. We found that while H1.2 or H1.5 OE had modest effects, only the combined H1.2+H1.5 OE drove a significant depletion of the open promoter signature (NES =-1.23, p = 0.081), indicating a synergistic role in chromatin compaction. In contrast, KD of these variants showed a non-significant profile with distinct enrichment scores and rank distributions compared to the OE conditions (Additional file 1: Fig. S7). These findings suggest that the specific, synergistic accumulation of H1.2 and H1.5 rather than their depletion is the primary driver of the’closed’ chromatin state observed in our 16p11.2 hemi-deletion models.

Our results imply that MAZ acts as a transcription suppressor of the examined H1 genes, and it is part of a regulatory network where MAZ associates with JUND and BACH1. This identifies MAZ haploinsufficiency as the primary molecular driver connecting the 16p11.2 hemi-deletion genotype to the pathogenic upregulation of specific linker histones. This mechanism reveals that H1 dosage is a central control for genome accessibility. Loss of MAZ in the 16p11.2 region disrupts this dosage, leading to a restricted chromatin state that likely interferes with brain development, as summarized in the mechanistic model shown in Fig. 6.

**Figure 6.**
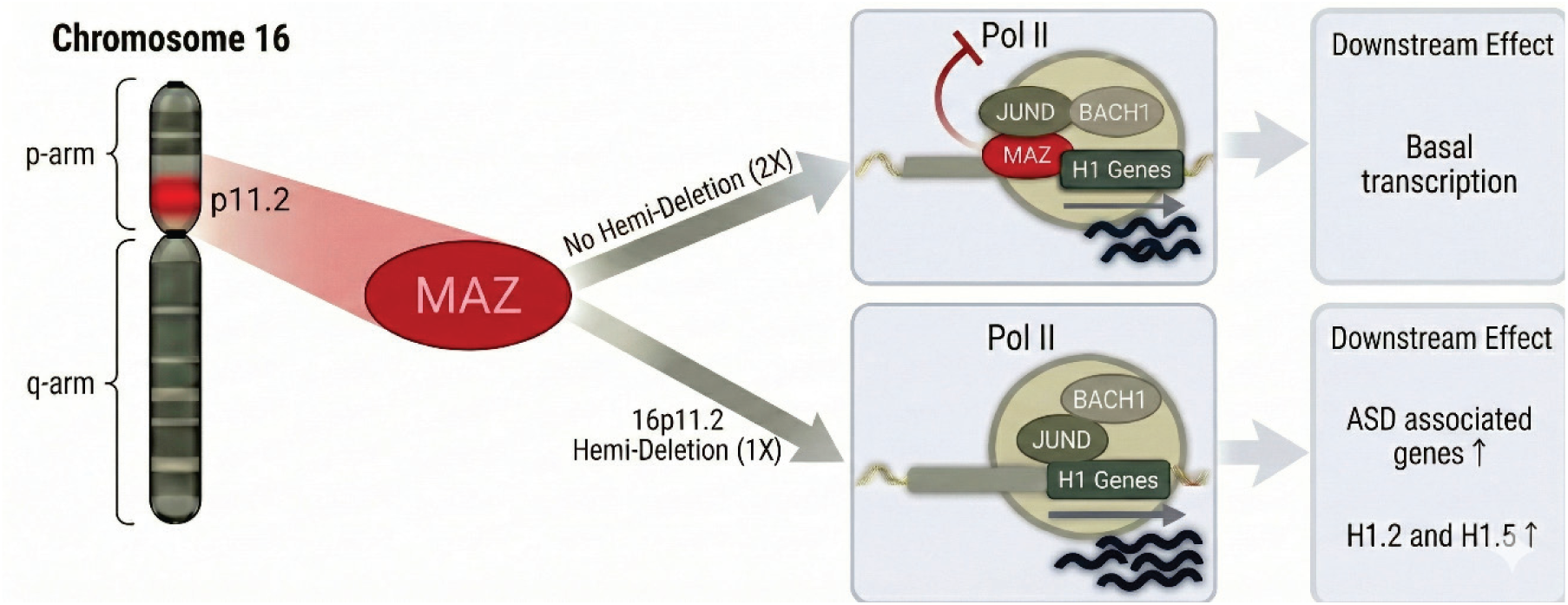
Mechanistic model of MAZ-mediated H1 regulation in 16p11.2 hemi-deletion. Schematic representation of the proposed regulatory mechanism linking 16p11.2 dosage to histone H1 dysregulation. (**Top**) Under normal conditions (two copies), MAZ binds GC-rich promoters of H1 genes and represses their transcription in cooperation with transcription factors such as JUND and BACH1, thereby maintaining basal transcriptional states. (**Bottom**) In 16p11.2 hemi-deletion (one copy), reduced MAZ dosage leads to derepression of H1 genes, particularly H1.2 and H1.5, resulting in elevated H1 levels. This increase in H1 dosage promotes chromatin compaction and alters downstream transcriptional programs, including dysregulation of ASD-associated genes.

## DISCUSSION

Our study reveals that histone H1 variants, particularly H1.2 and H1.5, are significantly upregulated in ASD with 16p11.2 hemi-deletion, but not in idiopathic SCZ or in BD, suggesting their selective involvement in specific neurodevelopmental pathways. This disorder-specific role in transcriptional regulation aligns with our prior studies indicating that both H1.2 and H1.5 regulate transcription through distinct chromatin occupancy patterns [15, 33]. Collectively, our results point to the involvement of H1 histones in early stages of neurodevelopment programs, specifically on glutamatergic synapse and translation regulation.

We identified the transcription factor MAZ as the primary mechanistic link driving H1 histone dysregulation. MAZ functions as a transcription suppressor of H1.2 and H1.5 expression, operating within a combinatorial hub alongside JUND and BACH1 to maintain proper repression. This regulatory control appears to be reinforced by higher-order chromatin architecture, as suggested by evidence of a physical interaction between the 16p11.2 locus and the HIST1 cluster [32], H1.2 and H1.5 might be positioned within regulatory loops that facilitate precise modulation by the MAZ complex. Consequently, 16p11.2 hemi-deletion depletes this repressive hub, causing an accumulation of linker histones. Increased H1 levels are known to tighten chromatin structure and lengthen the space between nucleosomes [51]. Our GSEA analysis confirms that genes with open promoters are repressed under these conditions. Therefore, we propose that MAZ haploinsufficiency causes’chromatin rigidity,’ which reduces the genomic flexibility needed for neurons to develop.

Network analysis revealed two major functional clusters affected by histone dysregulation: glutamatergic synapse function and 40S ribosomal subunit assembly. The histone-driven transcriptional changes in synaptic signaling pathways align with previous studies implicating excitatory synaptic dysfunction in ASD [34, 38]. Likewise, disruptions in ribosomal subunit assembly suggest that histone dysregulation may impact translational regulation, a process increasingly recognized as a key contributor to neurodevelopmental disorders [41, 42]. Beyond these neurodevelopmental pathways, our results also implicated broader cellular processes, including oxygen and hypoxia response, vasculature regulation, and apoptotic signaling, consistent with studies linking oxidative stress and metabolic dysregulation to ASD [52, 53]. Collectively, these findings support the view that linker histone dosage shapes neurodevelopmental gene regulation at a systems level. Beyond the locus-specific mechanisms described here, genome-wide, unsupervised analyses indicate that histone-driven transcriptional structure extends across ASD, SCZ, and 16p11.2 CNV contexts. This broader perspective suggests that altered H1 stoichiometry functions as a feature of disease-associated transcriptional programs rather than as an isolated, pathway-restricted effect.

Further supporting the importance of histone dosage in neurodevelopmental gene expression, we observed that complete knockout of H1.2 and H1.5 produced transcriptional profiles distinct from all other conditions, whereas overexpression and knockdown of both H1.2 and H1.5 clustered together resembled 16p11.2 CNV models. This finding highlights the need for precise dosage modulation: fully depleting histone variants does not simply reverse the effects of overexpression but instead induces an alternative transcriptional state. Such observations emphasize that balanced levels of H1.2 and H1.5, rather than their presence or absence alone, are pivotal to maintaining proper gene expression programs.

A striking aspect of our transcriptional analysis was the unexpected similarity in gene expression profiles among NPCs derived from individuals with 16p11.2 hemi-deletion and 16p11.2 duplication. We found that, despite their opposing genetic alterations, these conditions exhibited highly similar transcriptional signatures at early developmental stages, diverging only during neuronal differentiation. This observation suggests that histone dysregulation may act as an early-stage modifier of neurodevelopment, with later-stage differentiation amplifying condition-specific effects. This aligns with the neurodevelopmental hypothesis of psychiatric disorders [54], which posits that early transcriptional disruptions can drive distinct pathological trajectories depending on developmental timing and cellular context.

In conclusion, our study identifies the upregulation of linker histones H1.2 and H1.5 as a key molecular feature of 16p11.2-associated ASD. By integrating network analysis with functional validation, we map this dysregulation to the haploinsufficiency of the transcription factor MAZ, which normally represses the HIST1 cluster. When MAZ levels are low, the resulting imbalance in histone stoichiometry presumably leads to increased chromatin compaction. This structural change disrupts gene programs necessary for synaptic function and ribosomal assembly. Therefore, therapies designed to rebalance histone dosage or restore chromatin accessibility may provide new ways to address the deficits associated with this CNV.

## Conclusions

Our study defines a critical chromatin-mediated mechanism underlying 16p11.2-associated ASD. We identify that MAZ, a transcription factor located within the 16p11.2 region, binds to the promoters of H1 genes and acts as a transcriptional repressor. Due to the hemi-deletion of 16p11.2, the resulting reduction in MAZ levels leads to the upregulation of H1 transcription, particularly H1.2 and H1.5. We demonstrate that this specific dosage imbalance disrupts the genomic plasticity required for early neurodevelopment. Specifically, it impairs the transcriptional networks governing glutamatergic synapse formation and ribosomal assembly. These findings underscore that maintaining precise histone stoichiometry is fundamental for proper brain development and position the MAZ-H1 regulatory axis as a potential target for therapeutic strategies aimed at restoring chromatin accessibility in neurodevelopmental disorders.

### Methods Cell Culture

SH-SY5Y cells were cultured in DMEM/F12 medium (Merck), supplemented with 10% fetal bovine serum (Merck), 0.15% sodium bicarbonate, 2 mg/ml L-alanyl-L-glutamine, 100 U/ml penicillin, 0.1 mg/ml streptomycin (Sartorius). Cells were grown at 37 °C in a humidified atmosphere with 5% CO2.

### Overexpression Transfection

Transfection of the SH-SY5Y cell lines was performed by Lipofectamine 2000 reagent (Thermo Fisher) according to the manufacturer’s protocol. Three vectors were overexpressed: (a) pcDNA 4-TO-GFP, referred to as Mock, (b) pcDNA4-TO-H1.2, (c) pcDNA4-TO-H1.5. For each transfection, 6 million cells were seeded in 10-well plates, transfected with 5 μg plasmid, and incubated for 48 hours before total RNA extraction.

### Knockdown Transfection

Transfection of the SH-SY5Y cell lines was performed by Lipofectamine RNAiMAX (Thermo Fisher) according to the manufacturer’s protocol. Knockdown was performed using siRNA for H1.2 and H1.5 (ON-TARGET plus *HIST1H1C* and *HIST1H1B* Dharmacon’s smart pool, respectively, 100nM) and a control nontargeting SiRNA (D-001206 Dharmacon 100nM). 300-350K cells were seeded in 6-well plates one day before transfection and incubated for 72 hours before total RNA extraction.

### RNA isolation and cDNA formation

Total RNA extraction was performed using TRI reagent (Sigma) according to the manufacturer’s instructions. For RT-PCR reaction, 2 μg total RNA was reverse transcribed with SuperScript III First-Strand Synthesis System for RT-PCR (Invitrogen) with equal amounts of oligo(dT)20 and hexamers primers according to the manufacturer’s instructions.

### Quantitative RT-qPCR

The qPCR reactions were performed on an ABI StepOnePlus Real-Time PCR System (Applied Biosystems) using the following thermocycling parameters: 3 minutes at 95 °C followed by 40 cycles of 3 seconds at 95 °C and 30 seconds at 60 °C, ending with a dissociation curve. The calculation is based on the ΔΔCt method. Primer sequences are listed in the supplementary materials.

### RNA Sequencing

mRNA library preparation and subsequent sequencing were performed at Macrogen Europe (Amsterdam, Netherlands). 150-bp paired-end sequencing of mRNA samples from H1.2 OE, H1.5 OE, H1.2 + H1.5 OE, H1.2 KD, H1.5 KD, and H1.2 + H1.5 KD were performed using the Illumina NovaSeq 6000 system.

### Genome-Wide Association Study (GWAS) Gene-Based Analysis

Summary statistics from the most recent ASD genome-wide association study (GWAS) meta-analysis, publicly available through the Psychiatric Genomics Consortium (PGC) repository, were obtained. Specifically, we utilized the iPSYCH_PGC_ASD_Nov2017 dataset [1], which comprises summary statistics from a meta-analysis conducted by the Lundbeck Foundation Initiative for Integrative Psychiatric Research (iPSYCH) and the PGC, encompassing 18,381 ASD cases and 27,969 controls. For gene-based analysis, we employed PascalX [55], a Python3 implementation designed for high-precision gene and pathway scoring of GWAS summary statistics. PascalX implements the Pascal methodology for aggregating single-nucleotide polymorphism (SNP) p-values into gene and pathway scores. Using the default parameters of the genescorer class, we mapped individual SNPs to genes by extending gene boundaries by 50 kilobases (kb) upstream and downstream of transcription start and end sites, respectively. Given our focus on the HIST1 cluster, we restricted our analysis to chromosome 6. To visualize the gene-based association results, we generated a Manhattan plot using R (v4.5.2) and the ggplot2 package [56], focusing on a 4-megabase (Mb) region centered on the HIST1 cluster. The plot highlights genes achieving nominal significance (p < 0.05).

### HIST1 Gene Expression Analyses in Neurodevelopmental Disorders Using Publicly Available RNA-seq Datasets

Two independent studies were selected: (1) data from post-mortem brain samples of individuals with idiopathic ASD (n = 51), idiopathic SCZ (n = 559), and idiopathic bipolar disorder (BD, n = 222) [3], and (2) data from lymphoblastoid cell lines (LCLs) derived from 34 individuals from seven ASD-affected families carrying 16p11.2 CNVs [32]. The LCL dataset was included as a complementary, non-neural, patient-derived model in which the impact of 16p11.2 gene dosage on HIST1 expression could be assessed in a well-characterized familial cohort. Given the known cell type–specific differences in gene expression, LCL-derived results were analyzed separately and interpreted as supportive rather than primary evidence relative to brain and neural cell models.

The Gandal et al. dataset provided pre-computed differential gene and isoform expression summary statistics, including log_2_ fold changes and associated p-values, derived from their RNA-seq analysis of idiopathic samples compared to neurotypical controls. The Blumenthal et al. study reported genome-wide differential expression results, including standard log2 fold changes and permuted p-values derived from an ANOVA model incorporating genotype and sex, as well as beta coefficients derived from a reciprocal-effect linear model regressing expression on gene copy number. Differential expression results from both studies were filtered to retain only genes with the “HIST1” prefix in their gene symbols. Differentially expressed genes (DEGs) were defined as those with Benjamini-Hochberg (BH)-corrected p-values ≤ 0.05. To extend this analysis, fold-change values for HIST1 genes were compared with RNA-seq data from induced pluripotent stem cell (iPSC)-derived neuronal progenitor cells (NPCs) obtained from 10 individuals with 16p11.2 hemi-deletion, including both ASD and non-ASD subgroups [57], cerebral brain organoids generated from nine 16p11.2 hemi-deletion iPSCs [58], and NPCs derived from two individuals with 16p11.2 heterozygous duplication and SCZ [59]. For the 16p11.2 CNV models, differentially expressed genes were defined using Benjamini-Hochberg (BH)-corrected p-values < 0.05, with each CNV group (deletion or duplication) compared to its corresponding copy-number–matched control group.

An in-house R script was developed to visualize significant HIST1 gene expression changes using the ComplexHeatmap [60] and ggplot2 [56] packages. The final dataset included 27 HIST1 genes with expression changes, with 10 significant across all idiopathic disorders, 16 across all deletion carriers, 24 in duplication carriers, and 17 showing significant reciprocal HIST1 expression changes in 16p11.2 CNV ASD. Full details of differential expression analysis are provided in the supplementary methods.

### Normalized Gene Expression Analysis Across 16p11.2 CNV and Histone Models

Normalized gene expression data across multiple 16p11.2 CNV models were integrated from publicly available and newly generated RNA-seq datasets. Public datasets included RNA-seq data from neurons derived from three individuals with 16p11.2 duplication and SCZ [35], RNA-seq data from NPCs and cerebral organoids from individuals with 16p11.2 heterozygous deletions, with and without ASD [57, 58], and NPCs derived from individuals with 16p11.2 heterozygous duplication and SCZ [59]. To complement these datasets, we generated new RNA-seq data from neuroblastoma (SH-SY5Y) and embryonic kidney (HEK293) cell lines under different histone H1 perturbation conditions. In SH-SY5Y cells, H1.2 and H1.5 variants were overexpressed (OE) and knocked down (KD), while in HEK293 cells, somatic histone H1s were genetically knocked out (KO) [15]. For in-house SH-SY5Y perturbations, differential expression was assessed relative to matched controls (empty vector for overexpression and control non-target siRNA for knockdown), with two biological replicates per condition.

RNA-seq reads were aligned and quantified using VAST-TOOLS (v2.5.1) [61] with the --expr option, mapping to GRCh38. Gene expression was normalized using variance-stabilizing transformation (VST) in DESeq2 [62]. Principal component analysis (PCA) was performed using prcomp() in R, and Spearman correlations were computed on mean gene expression per condition, visualized with ggplot2 [56] and pheatmap [63].

To assess expression perturbation across 16p11.2 CNV and histone H1 models, we used the VST-normalized expression matrix and, for each gene, calculated the variability of expression across all conditions as the standard deviation of VST values. Genes were then ranked by this variability metric, which captures the magnitude of expression change irrespective of direction. For heatmap visualization, we selected the top 20 most perturbed genes, defined as those with the highest variability (no additional fold-change or p-value threshold was applied at this step, as this analysis was descriptive and rank-based).

Gene Ontology (GO) enrichment analysis of the top 500 perturbed genes was performed using clusterProfiler [64] with Benjamini-Hochberg (BH)-corrected p-values (FDR < 0.05), mapping gene symbols via biomaRt [65].

### Differentially Expressed Genes Intersections Across 16p11.2 and Histone Perturbation Models

DEGs were identified by comparing each experimental condition with its respective matched control. This analysis was performed across 16p11.2 hemi-deletion NPCs, cerebral organoids, NPCs and neurons from 16p11.2 duplication and SCZ cohorts, and histone H1 perturbation models (H1.2/H1.5 OE, KD, and histone H1 KO). For the UpSet plot, DEGs from all histone perturbation models were grouped into a single set, and DEGs from 16p11.2 hemi-deletion NPCs (ASD and non-ASD) were combined, allowing visualization of shared and unique DEGs across conditions using UpSetR [66].

PCA was performed on log_2_ fold-change values from specific DEG sets, where histone perturbation models (KO, KD, OE) and 16p11.2 hemi-deletion NPCs (ASD and non-ASD groups) were analyzed separately. The prcomp() function in R was used, and results were visualized using ggplot2 [56], with each condition color-coded.

### Differential Gene Expression Analysis of 16p11.2 Region Across Models

DEGs within the 16p11.2 chromosomal region were analyzed across all models. Histone perturbation conditions were initially grouped into OE, KD, and KO categories. For further analysis, only overexpression and knockdown conditions were expanded, while knockout conditions were omitted. Heatmaps were generated using ComplexHeatmap [60] with k-means clustering applied.

### Network Analysis of DEGs Across 16p11.2 CNV and Histone OE

DEGs from brain organoids with 16p11.2 hemi-deletion, neurons with 16p11.2 duplication and SCZ, and histone H1 overexpression (H1.2, H1.5, and combined) were analyzed using NetworkAnalyst [67]. DEGs with log₂ fold-change values were used to construct a generic protein–protein interaction (PPI) network, applying a minimum first-order interaction filter and restricting interactions to brain tissue.

Filtered network nodes were exported and analyzed using the STRING database [47], where k-means clustering was applied to identify functional gene clusters. log₂ fold-change expression profiles within each cluster were visualized using pheatmap [63] with hierarchical clustering.

### MAZ Regulatory Map, Promoter Binding and Knockdown Transcriptomes Analysis

Genes within the 16p11.2 interval (chr16: 29,638,676–30,188,531, GRCh38) were retrieved from Ensembl [68] and annotated. Transcription factor genes within this region were identified. Two transcription factors were identified, MAZ and TBX6. While MAZ is extensively studied, TBX6 is relatively unknown. In addition, MAZ has a known role as a GC-rich promoter–binding factor, which is found in the promoter region of all H1 genes. MAZ and its neighboring genes were then used to generate a MAZ-centered gene map. A MAZ-centered protein–protein interaction network was constructed in STRING [47], using MAZ as the seed and retrieving high-confidence interacting transcription factors under default parameters. Only experimentally supported and curated database interactions with a combined score ≥ 0.4 were retained, and the resulting network was exported and redrawn for presentation.

Promoter binding profiles for MAZ, JUND, and BACH1 at linker histone H1 genes were derived from publicly available ChIP-seq datasets in human cell lines (ENCODE [69]; accessions listed in Supplementary Table S2). Promoters of H1.1-H1.5 were defined as −4 kb to +0.5 kb around the annotated transcription start site (GRCh38). For each transcription factor and promoter, strand-aware per-base coverage was imported from bigWig files in R and averaged across biological replicates. Coverage vectors were oriented so that transcription always proceeded left-to-right, binned into 50 bp windows across the −4 kb to +0.5 kb interval, and smoothed using a 7-bin moving average to reduce local noise. To normalize for background, a baseline signal was calculated per transcription factor as the mean coverage in distal regions (−4 kb to −1 kb and +200 to +500 bp relative to the TSS), and the binned signal was expressed as fold over this background. Meta-profiles were plotted as transcription factor–specific average signal across position relative to the TSS using ggplot2 [56], with the TSS indicated by a vertical dashed line.

For transcription factor knockdown transcriptomics, we used RNA-seq datasets and expression arrays from siRNA experiments of MAZ, JUND, and BACH1 (GEO accession(s) listed in Supplementary Table S2). For each dataset, gene-level raw count matrices provided by the original studies were downloaded from GEO [70]. Samples transfected with TF-targeting siRNA were contrasted against matched nontargeting (control) siRNA. Counts were imported into R and analyzed using DESeq2 [62] with a design formula ∼ condition. Differential expression for each knockdown was assessed relative to the control, and p-values were adjusted using the BH method. Unless otherwise stated, genes with FDR < 0.05 were considered significantly differentially expressed. In parallel, log-transformed counts were analyzed with limma [71] to obtain moderated statistics and rank genes by differential expression. For visualization purposes in volcano plots, nominal p-value thresholds were applied as indicated in the corresponding figure legends. H1.1–H1.5 were specifically highlighted in volcano plots together with MAZ, JUND, and BACH1.

## Ethics approval and consent to participate

Not applicable.

## Consent for publication

Not applicable.

## DATA AVAILABILITY

The sequencing data generated in this study have been deposited in the Gene Expression Omnibus (GEO) under accession number GSE292323 and are publicly accessible at the GEO repository [72].

## Authors’ contributions

G.A. and R.B. conceived and designed the study. R.B. analyzed the genomic data. D.A., D.H., R.H., M.Z-O. and G.L.M. performed the RNA-seq experiments. D.A. and D.K. performed quantitative RT-PCR experiments. R.B. collected samples. G.A., I.K., and R.B. interpreted the results. G.A. and R.B. wrote the manuscript with contributions from all co-authors. G.A., G.L.M., M.Z., and R.E. supervised the project. All authors read and approved the final manuscript.

## ACKNOWLEDGEMENTS

R.B. is supported by a fellowship from the Edmond J. Safra Center for Bioinformatics at Tel-Aviv University. We also thank Macrogen Europe for their helpful work.

## Peer review information

Tim Sands was the primary editor of this article and managed its editorial process and peer review in collaboration with the rest of the editorial team. The peer-review history is available in the online version of this article.

## Additional files

Additional file 1. Supplementary Figures S1–S6, supplementary methods, primer sequences, and supplementary tables.

This file contains supplementary validation and transcriptomic analyses, detailed experimental and computational methods, qRT-PCR primer sequences, and tables listing antibodies and public datasets used in the study.

