## Supplementary material for "Histone H1 Variants Regulate Neurodevelopmental Transcriptional Programs in Autism with 16p11.2 deletion": Supplemtary file

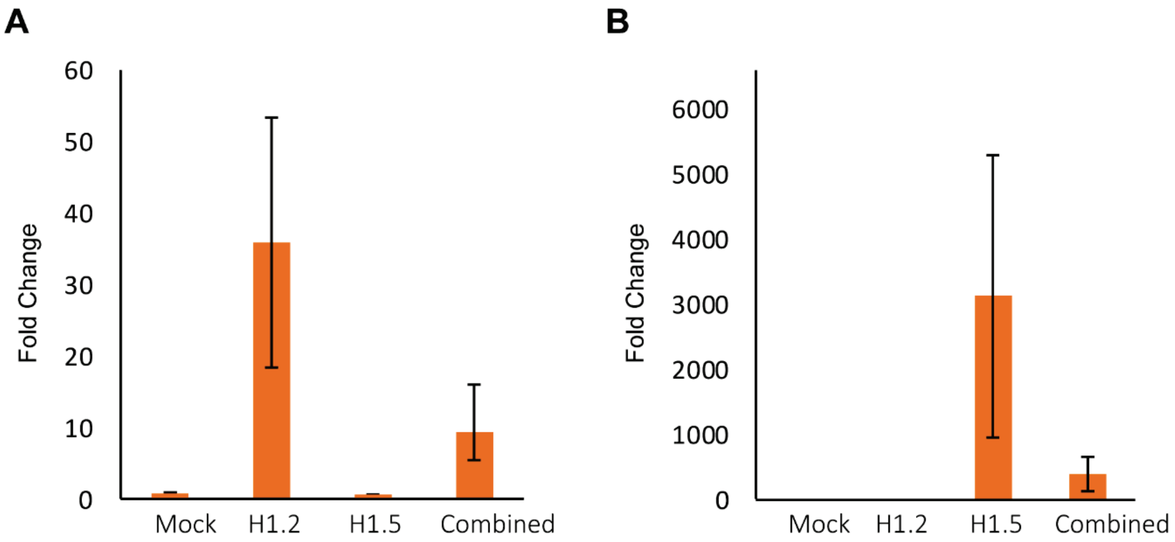

**Figure S1.** Validation of H1.2 and H1.5 overexpression. RT-qPCR analysis of histone variant expression 48h post-transfection. (**A**) H1.2 mRNA levels increased 35-fold following H1.2 transfection and 10-fold following co-transfection with H1.5. No significant change in H1.2 expression was observed in mock or H1.5-only transfected cells. (**B**) H1.5 mRNA levels increased 3000-fold following H1.5 transfection and 500-fold following co-transfection with H1.2. No significant change in H1.5 expression was observed in mock or H1.2-only transfected cells.

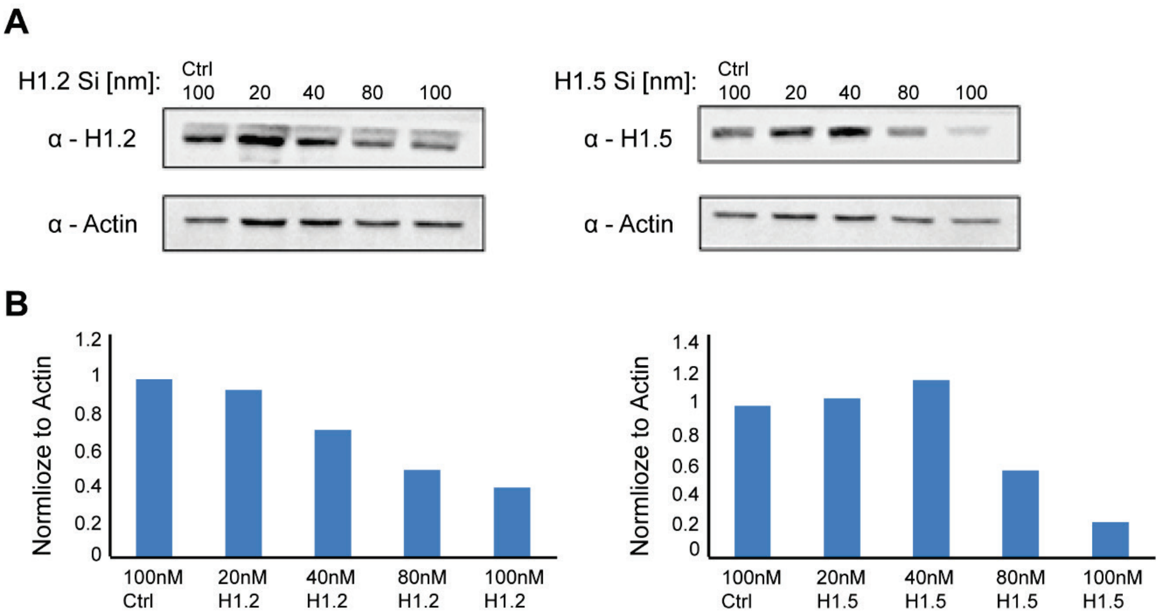

**Figure S2**. Calibration of the KD level. (**A**) SH-SY5Y cells were transfected with the indicated concentrations of siRNAs directed against H1.2 or H1.5 using Lipofectamine RNAiMAX. Whole-cell protein extracts were collected after treatment, and western blotting was performed with the indicated antibodies. Nontargeting siRNA was used as a control (Ctrl). (**B**) Quantification of western blot bands using ImageJ software. Values were normalized to actin levels.

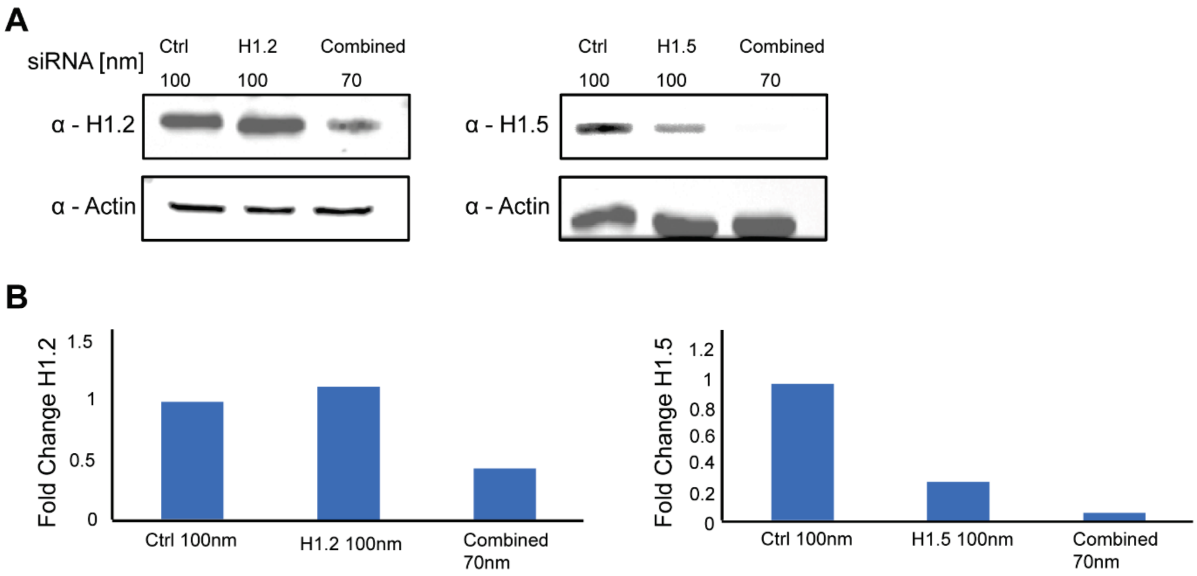

**Figure S3**. KD of H1.2, H1.5 and combined variants in SH-SY5Y cells: (**A**) The indication concentration of siRNA against H1.2, H1.5 and combined treatment was used to transfect SH-SY5Y cells, using lipoRNAiMAX transfection reagent. (**B**) WB analysis of total protein extraction was preformed, following 72h of transfection. ImageJ was used to quantify the bands. Values were normalized to actin bands.

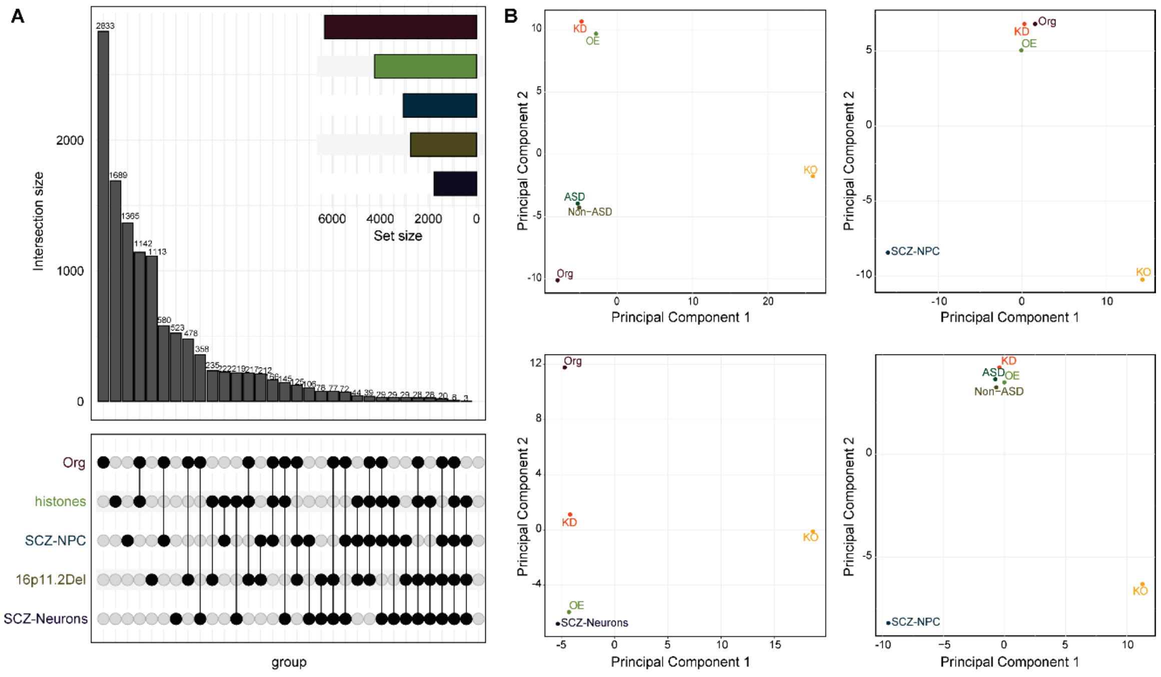

**Figure S4. Differential gene expression overlaps and principal component analysis across neurodevelopmental and histone perturbation models.** (**A**) UpSet plot of intersections between DEGs (condition vs. control) for five meta-groups: 16p11.2Del (NPCs from individuals with 16p11.2 hemi-deletion, with or without ASD), Org (cerebral organoids from 16p11.2 hemi-deletion iPSCs), SCZ-NPC (NPCs from individuals with 16p11.2 duplication and SCZ), SCZ-Neurons (neurons from individuals with 16p11.2 duplication and SCZ), and Histones (all H1.2/H1.5 OE, KD and KO conditions). For each meta-group, the union of DEGs across its constituent conditions was used. Vertical bars represent intersection sizes; horizontal bars denote total DEG counts per meta-group. (**B**) PCA plots based on log2 fold-change values for the union of DEGs across all conditions. Each point corresponds to an individual experimental condition; histone perturbation conditions are summarized into three groups (OE, KD, KO). Plots show the separation and proximity of histone perturbations relative to 16p11.2 hemi-deletion (ASD, Non-ASD and Org) and duplication (SCZ-NPC, SCZ-Neurons) models along the first two principal components.

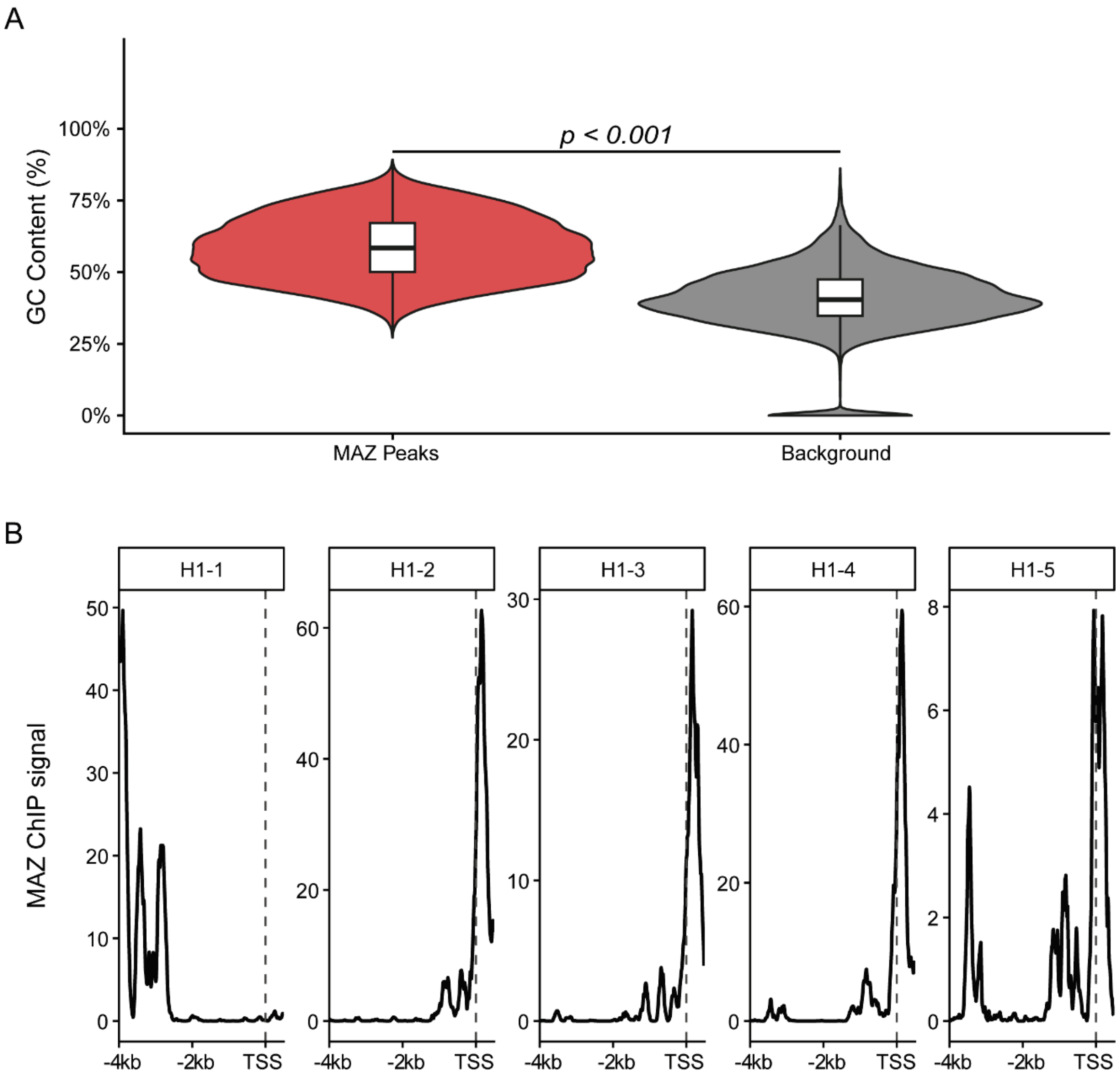

**Figure S5. GC-rich architecture and MAZ occupancy at H1-cluster promoters.** (**A**) (A) GC content analysis of MAZ binding sites. Violin plots compare the GC content percentage of MAZ binding peaks (red) against a random genomic background (grey). MAZ peaks exhibit significantly higher GC content (empirical $p < 0.001$, determined by permutation testing). (**B**) MAZ ChIP-seq signal profiles across extended promoter regions of individual H1 genes. MAZ ChIP-seq signal was extracted across −4 kb to +0.5 kb windows relative to the TSS for H1-1, H1-2, H1-3, H1-4, and H1-5. Signals were strand-aligned and plotted as normalized coverage; dashed vertical lines denote the TSS.

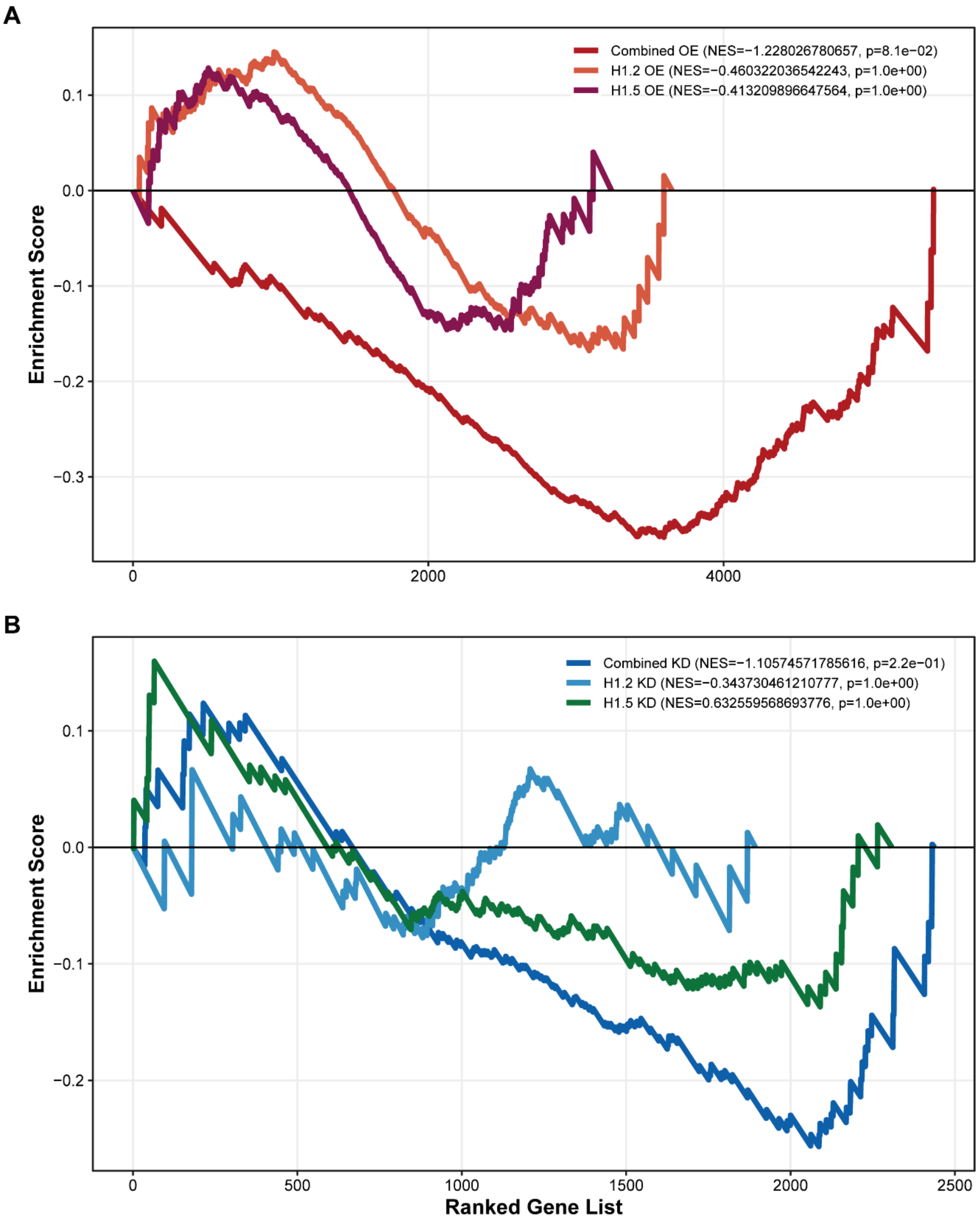

**Figure S6. Transcriptional response of high-confidence open chromatin genes to H1 dosage perturbation.** (**A**) GSEA enrichment profiles under H1 overexpression. The target gene set consists of high-confidence open promoters defined by re-analysis of basal SH-SY5Y ATAC-seq data (GSE274254); peaks were called using MACS2 (score > 1.0) and annotated to transcription start sites (TSS ± 3 kb). Genes were ranked by RNA-seq $log_2$ fold change for Combined H1.2/H1.5 overexpression (red), H1.2 overexpression (orange), and H1.5 overexpression (purple). Normalized Enrichment Scores (NES) and p-values (permutation test) display the degree of gene set downregulation. (**B**) GSEA enrichment profiles under H1 knockdown. Analysis performed as in (A), with genes ranked by $log_2$ fold change for Combined H1.2/H1.5 knockdown (red), H1.2 knockdown (orange), and H1.5 knockdown (purple) relative to scramble control. Curves reflect the distribution of the basal open chromatin signature across the ranked transcriptomes.

### Methods and Materials Extra:

#### Total protein extraction

Proteins extraction from cells pellets was done by a hypotonic lysis buffer (50 mM Tris-HCl, pH 7.5, 1% NP40, 150 mM NaCl, 0.1% SDS, 0.5% deoxycholic acid, 1 mM EDTA) containing complete protease inhibitor (Roche) and phosphatase inhibitor cocktails I and II (Sigma).

#### Western blot analysis

Proteins samples were separated by SDS-PAGE on 12% polyacrylamide gel and transferred by electro-blotting onto Protran nitrocellulose transfer membrane. The membranes were incubated with the appropriate primary and secondary antibodies and washed with TBS-Tween 20. Horseradish peroxidase conjugated secondary antibodies (Abcam). The antibodies and concentrations used are listed in Table S1 below.

| Target protein | Antibody catalog number | application | Western blot dilution |
|---|---|---|---|
| ACTIN | Millipore, MAB1501 | WB | 1:1000 |
| H1.2 | Abcam, Ab17677 | WB | 1:1000 |
| H1.5 | Abcam, Ab18208 | WB | 1:1000 |

#### Differential Expression Analysis

Sequencing reads were aligned and quantified to the human genome (Assembly hg38, GRCh38) using VAST-TOOLS (Irimia et al., 2014). VAST-TOOLS was run with default parameters and the --expr option to generate raw gene expression counts. Differential expression analysis was conducted using edgeR (Chen et al. 2025) through a custom R script. Low-abundance genes were removed using filterByExpr function with default thresholds. Library size normalization was applied using the trimmed mean of M-values (TMM) via the calcNormFactors function. Sample relationships were initially assessed through unsupervised clustering using both multidimensional scaling (MDS) plots and principal component analysis (PCA) with the plotMDS function applied to normalized counts. Batch effects, evident in both MDS and PCA plots, were corrected using ComBat-seq (Zhang et al. 2020), a batch adjustment tool specifically designed for RNA-seq count data. Batch adjustment was applied to the untransformed raw count matrix before subsequent analysis, using experimental batch as a covariate while preserving biological variation of interest. For differential expression testing, a negative binomial generalized linear model was fitted to the count data using the glmQLFit function with robust estimation. DEGs were identified using the quasi-likelihood F-test implemented in the glmQLFTest function. Genes were classified as differentially expressed if they met the dual criteria of false discovery rate (FDR) < 0.05 after Benjamini-Hochberg correction and p-value < 0.05. Log2 fold changes were calculated and reported for all DEGs.

**MAZ GC Content Analysis**

To assess the GC bias of MAZ binding, genomic sequences corresponding to MAZ ChIP-seq peaks were extracted. The GC content percentage was calculated for each peak and compared against a random genomic background generated by selecting random chromosomal segments of matching length. Statistical significance was assessed using a permutation test (n = 1,000 iterations) to derive an empirical p-value. Motif enrichment analysis was performed on peak sequences to validate the presence of the canonical MAZ binding signature (GGAGGG).

**Gene Set Enrichment Analysis (GSEA) of Open Promoters**

To evaluate the impact of H1 dosage on chromatin accessibility, Gene Set Enrichment Analysis (GSEA) was performed using the fgsea package in R. The target gene set consisted of high-confidence open promoters defined by re-analysis of basal SH-SY5Y ATAC-seq data (GSE274254). Accessibility peaks were called using MACS2 (score > 1.0) and annotated to transcription start sites (TSS ±3 kb) to identify genes with constitutively open promoter architecture.

For the enrichment analysis, genes were filtered to include only those identified as DEGs within the respective H1.2/H1.5 perturbation datasets. These DEGs were ranked by their average RNA-seq log2 fold change. Replicate gene entries were handled by averaging log2 fold change values prior to ranking to ensure a unique mapping between gene names and stats. Statistical significance was determined using the automated permutation algorithm in fgsea (seeded at 42 for reproducibility) to calculate Normalized Enrichment Scores (NES) and nominal p-values. Enrichment curves were generated to visualize the distribution of open promoter genes along the ranked lists, specifically comparing the synergistic effects of combined H1.2/H1.5 overexpression against individual variants and knockdown conditions.

**Primer sequences for qRT-PCR**

| Gene Name | Forward Primer | Reverse Primer |
|---|---|---|
| **SEZ6L2_485_ex12** | CCCTGGATCCAGGGATCTG | GTGACGGCTGTTGTCAGAGG |
| **SEZ6L2_932_ex34** | GTGACCAGCCCAGCCTAC | CGAATCACAGTGAAAGGTGGC |
| **KCTD13_000_ex1** | GACAGCAGAGACCCAATCAC | TCTCTGGTCTCGAGTCACC |
| **SNRPN (Outer)** | ACATCCACTGCAGCTGAGCT | CCGCGTGACATCTGCTTTT |
| **SNRPN (Inner)** | AGTCATTCTGCTTGCTGATC | CCTGAAGCTGTTAAGTGTTCTT |
| **DNMT1 (Outer)** | TGTTCTTCAGTTGGATTTAGGCC | CTCCCCGGTCTCCAGTC |
| **DNMT1 (Inner)** | ACAGCCCATCTTCCTGAC | AGGTCCCGCATGCAGGG |
| **GABRB3 (Outer)** | CCAACTTTGAATCTCCTGGCTT | AAAAATTTAATTCACCTGGGCAGT |
| **GABRB3 (Inner)** | ATCGAATCTTGGGGCAAAGAG | GAAAAATTTAATTCACCTGGGCAGT |
| **SRRM4_ex1** | GTCGCCAGTATCACGGCC | TTTCTTGTGCCTCTTCTCATCTC |

**Table S2**. Public datasets used for transcription factor binding and knockdown transcriptomic analyses (ENCODE and GEO accessions).

| **Factor** | **Data type** | **Source** | **Accession** | **Cell line / tissue** | **Assay / note** |
|---|---|---|---|---|---|
| MAZ | ChIP-seq | ENCODE | ENCSR163IUV | K562 | MAZ ChIP-seq in human cells |
| MAZ | siRNA RNA-seq | GEO | GSE215080 | K562, HAP1 | MAZ siRNA / KO RNA-seq (MAZ depletion vs control) |
| JUND | ChIP-seq | ENCODE | ENCSR000EGN | K562 | JUND ChIP-seq in human cells |
| JUND | Expression profiling by array | GEO | GSE86193 | HEKn | JUND SiRNA |
| BACH1 | ChIP-seq | ENCODE | ENCSR000EGD | K562 | BACH1 ChIP-seq in human cells |
| BACH1 | siRNA RNA-seq | GEO | GSE165718 | Calu3 | BACH1 SiRNA |
